# Dynamic Hh signaling can generate temporal information during tissue patterning

**DOI:** 10.1101/451872

**Authors:** Diana García-Morales, Tomás Navarro, Antonella Iannini, David G. Míguez, Fernando Casares

## Abstract

The differentiation of tissues and organs requires that cells exchange information in space and time. Spatial information is often conveyed by morphogens, molecules that disperse across receiving cells generating signaling gradients. Cells translate such concentration gradients into space-dependent patterns of gene expression and cellular behavior [1, 2]. But could morphogen gradients also convey developmental time? Here, investigating the developmental role of Hh on a component of the *Drosophila* visual system, the ocellar retina, we discovered that ocellar cells use the non-linear gradient of Hh as a temporal cue, collectively performing the biological equivalent of a mathematical logarithmic transformation. In this way, a morphogen diffusing from a non-moving source is decoded as a wave of differentiating photoreceptors that travels at constant speed throughout the retinal epithelium.

## RESULTS AND DISCUSSION

Morphogens of the *hedgehog(hh)*/Shh family contribute spatial information during the development of a wide range of organs and organisms [3]. In addition, during the development of the *Drosophila* compound eye, Hh drives a wave of photoreceptor (R) cell differentiation across the eye primordium at a constant speed [4, 5]. A similar Shh moving wave has been described during the differentiation of the ganglion cells in the zebrafish retina [6]. However, these waves are not generated based on the morphogen character of Hh/Shh (i.e. differential responses to varying Hh concentration in space), but on the fact that the source of Hh production itself moves across the developing retina: Hh/Shh molecules non-autonomously induce progenitors to differentiate into retina cells which, in turn, start producing Hh/Shh. In this way, the source of signaling molecule moves coupled to the differentiation process [5, 6].

In addition to the compound eye, Hh signaling is necessary for the specification and differentiation of the *Drosophila* ocelli [7–9], three small eyes (one anterior and two posterior) located on the fly’s forehead that are part of the visual system of most insects (Figure 1A,B). Ocellar differentiation takes place in the dorsal-anterior region of the eye-antennal imaginal disc (Figure 1B). Here, one domain of Hh expression is flanked by two regions competent to differentiate into the ocellar photoreceptors (R cells) under the action of Hh signaling. One marker of competence is the gene *eyes absent (eya)*([8, 9])(Figure 1C,D). When the two contralateral discs fuse, the anterior ocellar regions merge into a single anterior ocellus (aOC), while the two other regions remain separate and will develop into the paired posterior ocelli (POC). Here we focused on the larger posterior ocellus to study how ocellar progenitor cells differentiate. R differentiation can be followed using the neuronal markers Elav and Glass. We observed that R cell differentiation proceeded in a wave-like fashion –i.e., differentiation starts in the vicinity of the Hh source, to then progress in the proximal-distal direction across the ocellar tissue (Figure1E). The transition from precursors to R cells can be monitored using the precursor marker gene *senseless (sens)*([10]) (Figure 1-Figure S1A-C). We find that, like in the compound eye, Sens expression precedes temporally that of Elav. Sens expression in differentiating R cells is transient, and it decreases as Elav expression increases. Spatially, Sens and Elav distribute along a proximal-distal axis with respect to the Hh source. Therefore, the differentiation wave can be visualized as a succession of Elav and Sens along this axis, with new Sens expressing cells being added progressively further away from the Hh source as differentiating cells express Elav and downregulate Sens (Figure 1-Figure S1). Importantly, and in contrast to the moving wave of Hh that sweeps across the developing compound eye, Hh is never expressed in ocellar R cells (Figure 1-Figure S1D,D’)[9, 11]. The Hh source remains the inter-ocellar region and therefore, does not move in space. To start investigating the potential role of Hh signaling in organizing this wave, we first examined the distribution of Hh across the competence domain, which is about 40μm (10 cells) wide, using a Hh:GFP BAC construct [12]. Hh:GFP disperses away from its source following roughly a decaying exponential, that can be fitted in space and time using a polynomial function (Figure 2A,A’ and see below). The Hh receptor Patched (Ptc) is also a target of the signaling pathway, so that its expression can be used as a read-out of the pathway’s signaling activity [13]. We found that, before R differentiation starts, Ptc expression follows the Hh:GFP gradient (Figure 2A,A’ and see below), indicating that signaling intensity reflects Hh distribution across the ocellus. In addition, this result suggested that the non-uniform Hh distribution could contribute to generating the wave, transforming the spatial gradient into a temporal axis, such that cells closer to the Hh source (and therefore, receiving higher concentration of Hh) would differentiate earlier than cells farther away. To test this possibility, we equalized Hh signaling across the developing ocellus by expressing, specifically in the ocellar primordia, uniform levels of *cubitus interuptus (ci)*, the Gli-type nuclear transducer of the Hh pathway [14], (Figure 2B,C and Figure 2-Figure S1A,B). After Ci overexpression, a larger than normal number of cells had initiated the expression of Sens and Elav relative to control ocelli, indicating their premature differentiation. More importantly, the progression of the wave was disrupted: instead of the succession of Elav and Sens cells, in Ci-overexpressing ocelli Elav and Sens cells are intermingled (Figure 2B,C). This result was compatible with the idea that the Hh signaling gradient encodes a temporal axis that generates the wave-like differentiation of ocellar R cells. To test this point more directly, we distorted the normal distribution of Hh by inducing new foci of Hh expression from around the developing ocelli (*wg2.11*-GAL4; *UAS-GFP:Hh* or “*wg>Hh*”; [15] and Figure 3) to then compare the spatial patters of Elav+Sens-, Elav+Sens+ and Elav-Sens+ cells between control and *wg>Hh* ocelli (Figure 3A-D’). Since even the wild type pattern shows some variability, we used a statistical analysis to compare the “grouping” (as measured by the departure from a random proportion of neighbors of a given type) and “polarity”, which measures the ordered succession of cells states along the proximodistal axis (and that is a defining trait of a wave) (Figure 3E-G’), of these patterns. Control and *wg>Hh* patterns were both significantly-but similarly-different from random (Figure 3 S1A), as expected if spatially localized Hh drives the pattern of differentiation. However, when “polarity”, the statistic that reflects a wave-like organization, was analyzed, control samples were significantly more polarized than *wg>Hh*, which were closer to a non-polarized distribution (Figure 3 S1B; see Methods for a complete description of the statistical analysis). These results confirm that, despite the variability of the system, the pattern of differentiation from Sens precursors to Elav photoreceptors progresses as a wave, and reinforces the notion that the distribution of Hh across the developing ocellus is necessary for organizing this wave. Next, we tested whether blocking Hh signaling could result in abrogation of R differentiation. To do that, we expressed a dominant negative Ptc receptor (PtcΔloop2 which, due to its incapacity to bind Hh, represses the pathway constitutively [2]). Our results show that, as in the compound eye, Hh is necessary for R differentiation in the ocelli (Figure 2-Figure S1C-E). Altogether, our results so far indicated that the time needed for a cell to start differentiating depends on the amount of Hh that it receives. Because Hh distribution decays non-linearly in space (Figure 2A’), R cells should also accumulate non-linearly over time (i.e. with fast R generation close to the source and progressively slowing down with increasing distance from it). To test this hypothesis, we quantified the number of Elav-expressing R cells over developmental time. As developmental timer we used the number of rows of ommatidia that have undergone differentiation in the compound eye, which is known to increase at a constant speed [16, 17]. In contrast with the expectation, the number of Elav cells increased linearly with time in both anterior and posterior ocelli, indicating that the differentiation wave propagated at a constant speed (Figure 2D).

**Figure 1.**
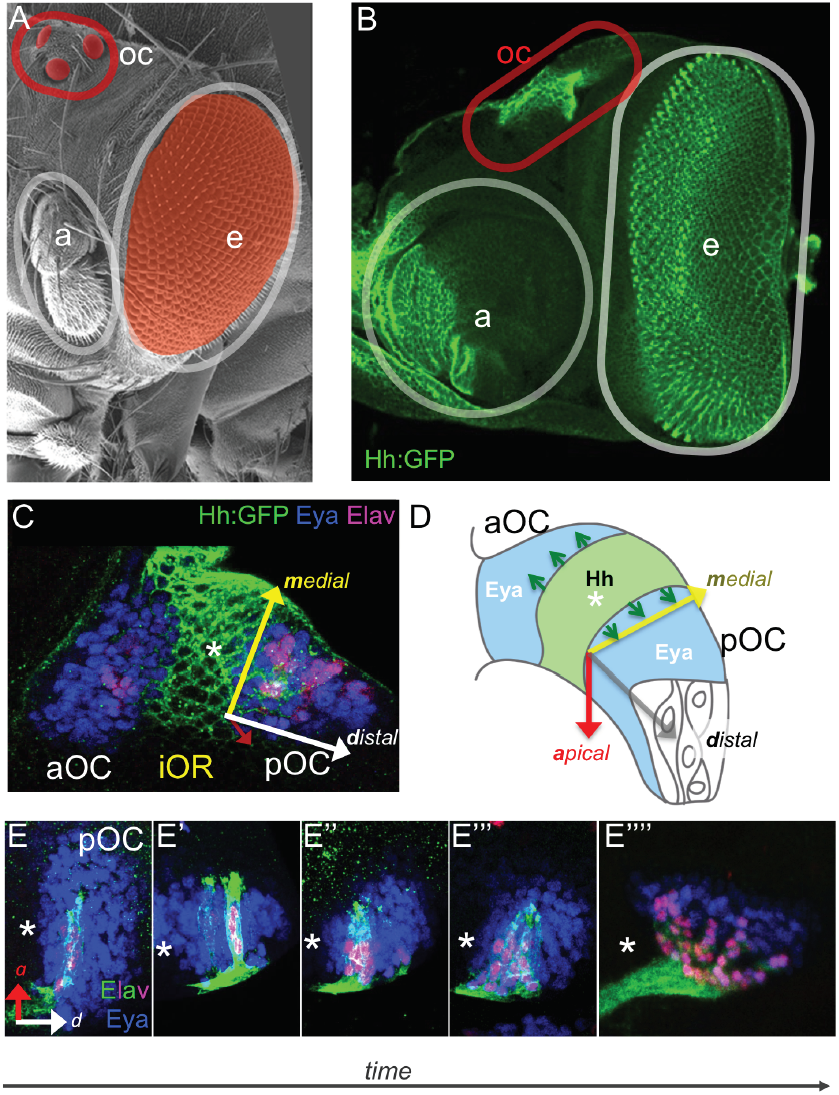
Photoreceptor (R) differentiation in the Drosphila ocelli. (A) SEM view of a *Drosophila* head. Outlined, the ocelli (“oc”), the compound eye (“ce”)–both pseudocolored) and the antenna (“a”). (B) Confocal image of an eye-antennal head primordium of a Hh:GFP-BAC larva (late third instar) marking the prospective “oc”, “ce” and “a”. Hh:GFP in green. (C) Close up of the prospective ocellar region of a Hh:GFP-BAC primordium (green) stained for Eya (competence marker, blue) and Elav (neural marker, magenta). Hh is produced from a central domain that will become the interocellar region (iOR). The position of the Hh-expressing domain is marked by the asterisk (*) in C-E’’’’. Adjacent to it, the anterior and posterior domains of Eya-expressing cells will become the anterior (aOC) and posterior (pOC) ocelli, respectively. (D) Schematic representation of the ocellar region, showing the Hh-producing and Eya-expressing domains. The arrows indicate the spatial axes. (E-E’’’’) Temporal series of pOC regions from progressively older larvae/early pupa (as indicated by the “time” arrow), marked with Eya (blue) and Elav (Elav>nRFP_ires_mGFP). Images are from different, fixed discs. Elav-expressing photoreceptor (“R”) cells appear first closest to the Hh source (E) and then accumulate successively in more distal regions (E’-E’’’’). Nuclei and membranes of Elav cells are marked in magenta and green, respectively.

**Figure 2.**
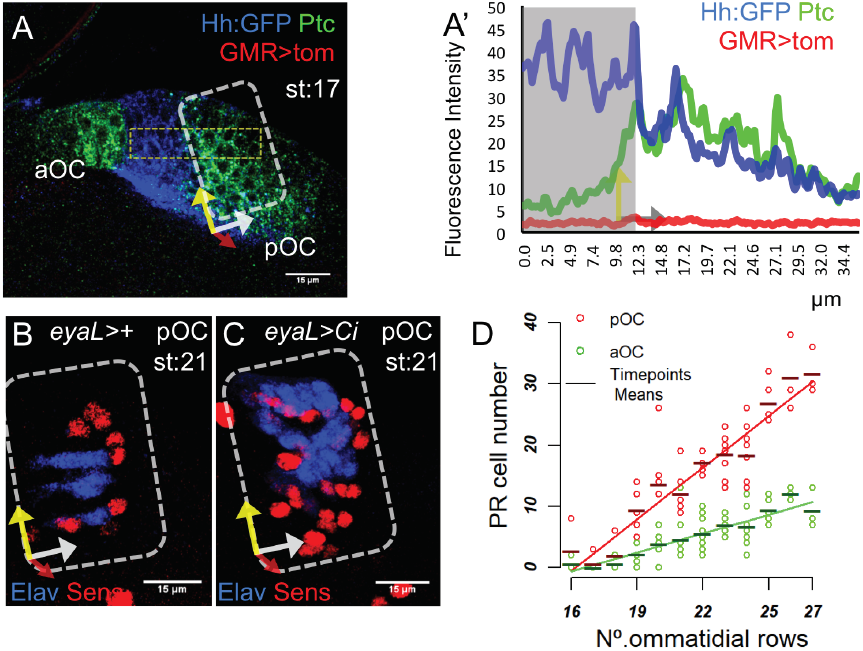
Hh signaling and R differentiation wave. (A) Confocal image of the ocellar region of a Hh:GFP; GMR>tdTomato (“GMR>tom”) larva (stage 17 ommatidia), stained for Hh:GFP (blue), Ptc (green) and anti-Tomato (red). No R cells (“GMR>tom”) have as yet differentiated. (A’) Quantitative profiles of the Hh:GFP, Ptc and GMR signals across the Hh producing domain (shaded in grey) and the pOC (measured in the dashed yellow box in (A)). Hh:GFP signal decays non-linearly. Ptc signal follows that of Hh:GFP at this stage, when no R cell (GMR>Tom) has differentiated yet. (B,C) pOC regions (boxed, like the corresponding region in (A)) stained for Elav (red) and Sens (blue) of discs from larvae of the same stage (21 ommatidia). In the control (B, “eyaL>+”) a row of R-expressing Elav cells precede a row of Sens-expressing precursors. In eyaL>Ci (C, causing the uniform and strong expression of Ci) precocious differentiation is observed. In addition, the differentiation wave, characterized by the succession Elav→Sens, is broken. (D) Number of Elav-positive cells in the pOC (red) and aOC (green) as a function of developmental time. The number of ommatidial rows in the compound eye, which increases linearly with time, was used as internal developmental timer. Data (circles) and means (“—”) are represented and fit well to a line. See Methods for a description of the statistical analysis. Source data for Figure 2 A’, D available as supplementary material (SD_Figure2_A’_D).

**Figure 3.**
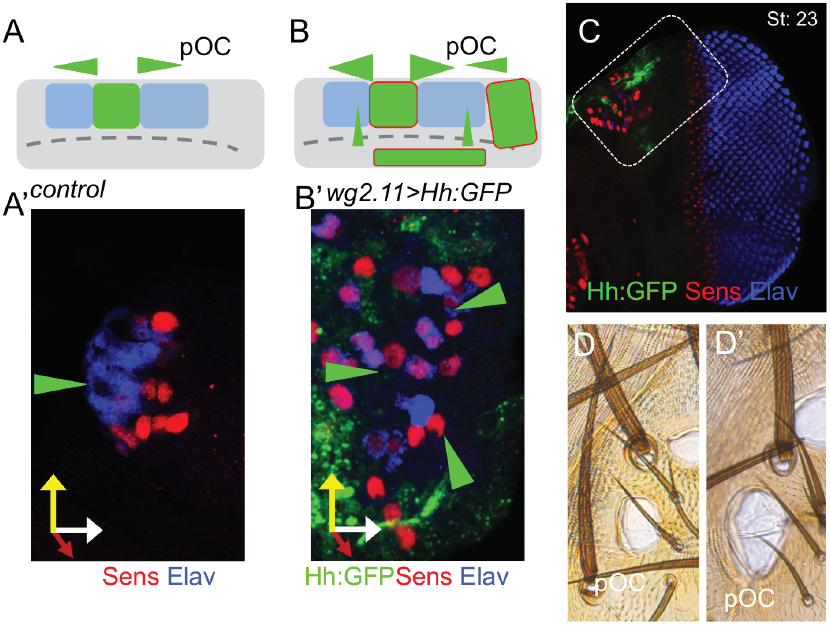
Altering Hh spatial distribution distorts the differentiation wave. (A,B) Cartoon depiction of the Hh sources (green domains) relative to the retina competent regions (blue) in control (A) and *wg>Hh* (B) ocellar regions. The posterior ocellus is marked as “pOC”. The green triangles indicate the distribution of Hh from these sources. In *wg>Hh*, Hh is expressed around the ocelli and within the normal Hh expression domain. (C) Late *wg>Hh* disc (st:23) stained for GFP (GFP:Hh), Sens and Elav. The boxed region corresponds to that represented in A and B. A’ and B’ are pOC regions from control and *wg>Hh* individuals, respectively. (D,D’) ocelli of control and *wg>Hh* adults. In *wg>Hh* ocelli are larger.

In order to explore the signaling outputs in this system, we constructed a mathematical model capturing the essence of the Hh signaling pathway (see Methods). In this model, Ptc represses Hh signaling targets unless it binds Hh. As Ptc is one of the pathway’s targets, Hh binding to Ptc results in the releases Ptc’s repressive action and results in its upregulation [13]. Sens is included as a target in the model, although this does not imply that Sens is a *direct* target. Expression of Elav in Sens-expressing precursors follows irreversibly, and to reflect the loss of Sens expression observed in Elav cells, we have also included a negative feedback from Elav to Sens (see Figure 4A). The dynamics of Hh production and dispersion in the model were calibrated using measured Hh:GFP profiles determined experimentally (Figure 4-FigureS1; and Figure 4-STable). The intrinsic variability of the system is modeled by introducing a 20% variability in all parameters of the model (see Methods). With no further assumptions, the model simulations confirmed the prior expectation: the accumulation of Elav cells was non-linear and differentiation was often not completed during the developmental period allowed (40 hours)(Figure 4B). The fact that the model was unable to reproduce the experimental observations indicated that our understanding of the signaling dynamics was missing some important process. Due to the key relevance of Ptc as both Hh receptor and target of the pathway, we examined in detail the dynamics of Ptc accumulation during differentiation. We found that, while before R cell differentiation Ptc signal followed a non-linear decay with a strong peak close to the Hh source (see Figure 1A,A’), in later stages Ptc signal decreased dramatically in R cells, identified by expression of Elav (Figure 4C-D’). Since binding of Hh to Ptc reduces its mobility [18], we reasoned that the reduction of Ptc availability in R cells could allow the non-bound Hh to move over these cells and disperse farther away from the source. In this model, the sequential dampening of Ptc expression acts as a “desensitization” mechanism. When this Ptc dampening was incorporated in the model (simplified as a repressor link from Elav-R to Ptc) it now correctly predicted that the differentiation wave moves with about constant velocity, and achieves the full differentiation of the progenitor population during the differentiation window (Figure 4E) (See also Figure 4-Supplementary Videos 1 and 2). In addition, the simulated dynamic profile of Hh matched that measured experimentally (Figure 4-Figure S1), with its gradient flattening and reaching further as developmental time progresses. Simulations include the approximate 50% reduction of Hh:GFP production that we observed experimentally (Figure 4-Figure S1A) but the results remain the same if the Hh production rate is maintained constant (Figure 4-Figure S2). Therefore, the desensitization of differentiating cells to Hh, caused by the dampening of Ptc, would allow the field of ocellar competent cells to transform the Hh gradient into a moving signaling and differentiation wave of constant speed. It has been described that Ptc is down-regulated upon binding to Hh [19–21] and also in a self-regulated manner [22] in *Drosophila* wing discs. To test if the dramatic downregulation of Ptc we observe was due to R cell differentiation or just to a process depending on ligand binding or Ptc concentration, we examined Ptc levels in the ocelli of late discs from *atonal (ato)* mutant larvae, in which R differentiation is abrogated (Figure 4-Figure S3). For each disc, the levels of Ptc signal in the ocelli were normalized relative to the signal in the antenna of the same disc. While in control discs *(ato1-/*+) the relative levels of Ptc decrease with time (Figure 4-Figure S3A,B), Ptc expression is maintained at levels comparable to those found in control discs before R differentiation onset, despite their having been exposed to Hh for the whole duration of the third larval stage (Figure 4-Figure S3C). To test directly whether R differentiation was causing Ptc downregulation, we drove uniform and premature Sens expression to force ocellar cells to differentiate prematurely. As expected after Sens overexpression, the ocellar region of *eyaL>Sens* larvae had an increased number of Elav-expressing cells (relative to stage-matched controls). These cells also showed a concomitant loss of Ptc expression (Figure 4-Figure S3F,G). Therefore, in the ocelli, R differentiation is a major controller of Ptc dynamics.

**Figure 4.**
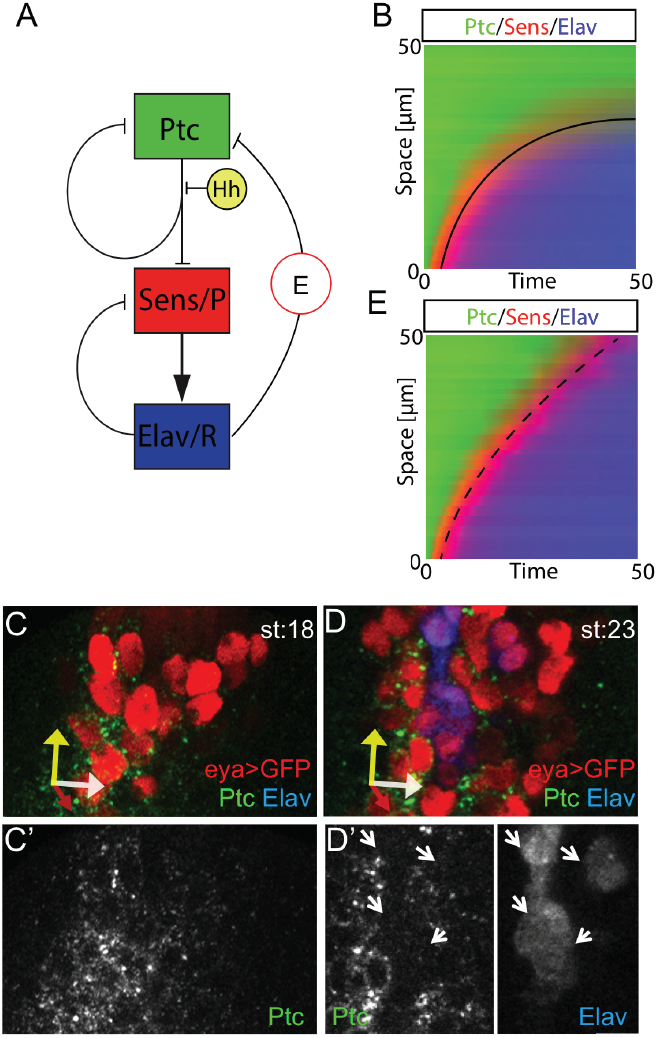
Loss of Ptc in R cells suffices to explain linear differentiation dynamics. (A) Cartoon diagram of the model for the Hh signaling pathway and its downstream effects. (B and E) Spatio-temporal dynamics of the model’s outputs without considering (B) or considering (E) a negative feedback from Elav-expressing R cells to Ptc (“E” link in (A)). In (B) R cells (blue) accumulate hyperbolically and do not reach the end of the competent region within the time frame of 50h. In (E) (with negative feedback, all other parameters being the same) R accumulation dynamics is close to linear and R differentiation reaches the end of the competent region. Simulations have been carried out including a 50% reduction in Hh production rate along the 50 h time, as observed experimentally. Similar results are obtained if this rate is maintained constant (Figure 4-S2). (C,D) Ocellar region of a st:18 (C,C’) and st:23 (D,D’) *eyaL>GFP* disc, stained for GFP (marking the Eya-expressing competence domain), Ptc and Elav (R cells). Axes as in Figure 1. Elav-expressing cells (marked by arrows) show reduced levels of Ptc. Source code for Figure 4 is available as supplementary material (SD_Figure4_script_Hhpathwaymodel).

One important aspect of ocellar differentiation is that, by the end of development, the number of R cells per ocellus is very consistent (the number of R cells of the adult posterior ocellus is 47,9; s.d.=0,7; n=5). However, we have noticed in static measurements of Hh:GFP that its signal is highly variable. To test for robustness, we compared the output of the model with or without signal desensitization, varying Hh production rates up to 10%. While without desensitization dynamics were far from linear, and the time to differentiation termination varied widely, the model including reduced Ptc availability coupled with differentiation maintained linearity (i.e. constant differentiation speed), and showed low variability in time to termination, despite these variations in production rates (Figure 5 and Figure-FigureS1). Therefore, our model predicts that, in addition to promoting a differentiation wave of constant speed, the R differentiation-induced Ptc desensitization results in increased developmental reliability.

**Fig 5.**
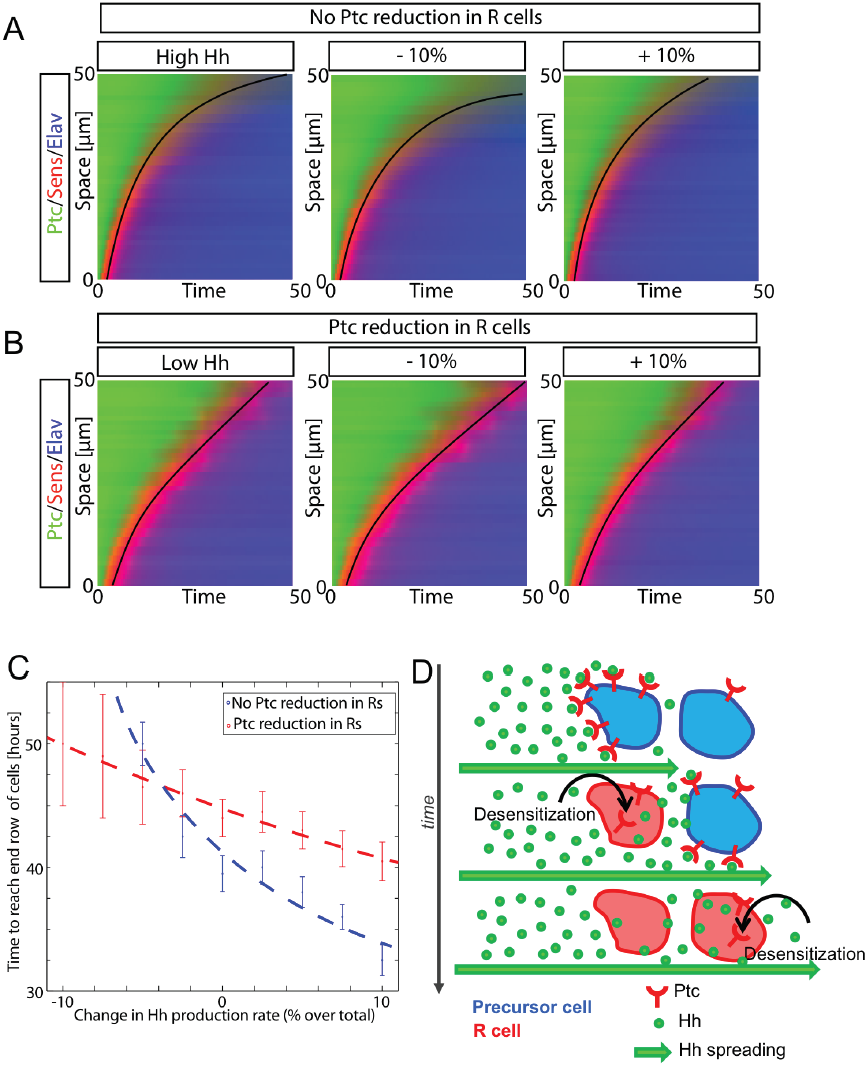
Robustness in the dynamics of the wave against changes in Hh increases when Ptc feedback is present. (A,B) Spacetime plots when Hh concentration is increased and decreased by 10% compared in the absence (A) or presence (B) of Ptc reduction in R cells (B). Colors represent the expression levels of Ptc (green), Sens (red) and Elav (blue). The solid line is used to represent the speed of the wave as a guide to the eye. The intensity of Hh in the left panel in (A) has been adjusted to facilitate comparison between (A) and (C). (C) Changes in the dynamics of the wave due to changes in Hh concentration. The model with no Ptc reduction (blue dashed line) is more sensitive to changes in Hh concentration that the situations with Ptc reduction. Statistics performed using 30 independent simulations for each point. Bars correspond to the standard deviation of each measurement. (D) Schematic depiction of the model proposed. Hh spreading leads to Ptc upregulation and maximal signal first closest to the source. As the cells differentiate, Ptc levels decrease allowing farther extension of Hh spreading. By each cell dynamically responding to Hh, the ocellar primordium transforms a noisy, non-linearly decaying signal into a differentiation wave of constant speed that is robust to signal noise. Source code for Hh signaling model available as supplementary material (SD_Figure4_script_Hhpathwaymodel).

Previous work has shown, in different developmental contexts, how a spatially static source of Hh/Shh coupled to its dynamic intracellular signaling network can generate spatial patterns of gene expression [23, 24]. In this paper, we show that a similarly static Hh source can be decoded as a linear “time arrow” – a wave of differentiation of photoreceptors with constant speed. This capacity requires a single change in the regulation of the Hh receptor Ptc. Two system-level properties are worth mentioning: First, the “log-transform” of the gradient’s signal is an active process, in the sense that cells are not passive readers, but transform the signal dynamically through a reactive intracellular signaling network. Second, the mathematical transformation of the signal is an emerging property of the system: While the signaling changes operate at a single cell level, this transformation requires a number of cells coupled within a Hh gradient. Even though the pervasive use of Hh/Shh as a morphogen might be the result of evolutionary contingencies, an alternative explanation is that Hh and its signaling pathway, acting on fields of cells, is flexible in the type of information outputs cells generate when reading the gradient. It is conceivable that this flexibility would be a selective advantage that might have resulted in the Hh signaling pathway being redeployed once and again during evolution.

## ACKNOWLEDGEMENTS

We thank T. Kornberg (UCSF), C. Sánchez-Higueras and J CG. Hombría (CABD), G. Struhl (Columbia Univ.) and J. Culí (CBMSO) for fly strains; A. Jarman (Univ. of Edinburgh), R. Holmgren (Northwestern Univ.), B. Hassan (ICM), R. Barrio (Biogune), X. Franch (IBE-UPF) and I. Guerrero (CBMSO) for antibodies; and the CABD ALMI platform for imaging and image analysis support.

## Authors’ contributions

FC conceived the study, acquired funding and managed the project; FC, DG-M and DGM designed research; DG-M carried out experimental work with the help of AI during the revision phase; TN devised the statistical tests and carried out statistical analyses; DGM acquired funding and developed model; FC, DG-M and DGM analyzed experimental and modeling results, FC drafted the manuscript and all authors contributed to the final manuscript.

## Data and Material availability

all data is available in the manuscript or in the supplementary materials.

---

**Figure 1-Figure S1.**
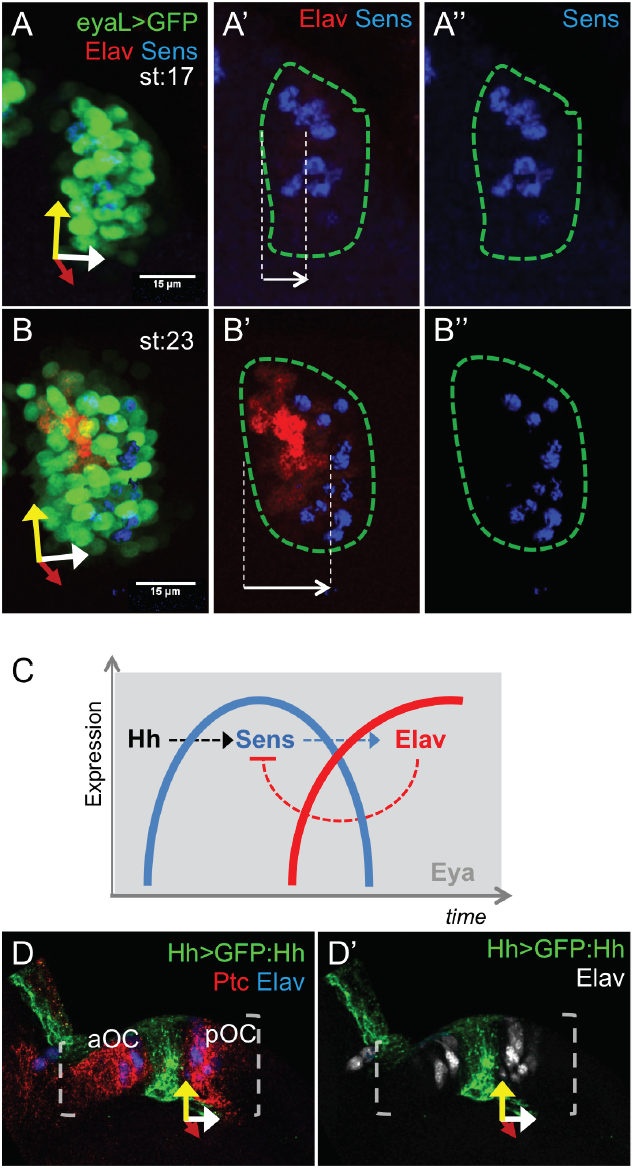
R cells do not express Hh and differentiate following a *senseless-Elav* sequence. (A,B) Sens and Elav mark the progress from precursors (Sens, red) to differentiating photoreceptors (Elav, blue). Confocal image of a posterior ocellar region of third instar stage 17 ommatidia (A) and 23 ommatidia (B) discs of an *eya>GFP* larvae. Eya (green) expression marks the ocellar competent region (outlined in A’-A’’ and B’-B’’). At St17 Sens is expressed adjacent or close to the proximal border of the ocellus and no Elva R cells have yet differentiated. At St23, Elav R cells have differentiated and new Sens-positive cells are induced distal to them. The white arrows in A’ and B’ indicate the position of the Sens front relative to the Hh source. Axes as in Figure 1. (C) Schematic representation of the temporal changes in gene expression experienced by any cell in the ocellar complex. Competent cells (expressing Eya), upon receiving Hh signal, progress along their differentiation program, expressing Sens first, then Elav. The connecting links are dashed to indicate that the activations (arrow) or repression (flat end) are likely indirect. (D,D’) Confocal image of an ocellar region (bracketed) from a *hh-Gal4>UAS-GFP:Hh* disc, stained for GFP, Ptc and Elav (D). (D’) shows the GFP and Elav channels only. Elav-expressing R cells, which differentiate in a region of Hh signaling (i.e. Ptc-expressing), do not transcribe Hh.

**Figure 2-Figure S1.**
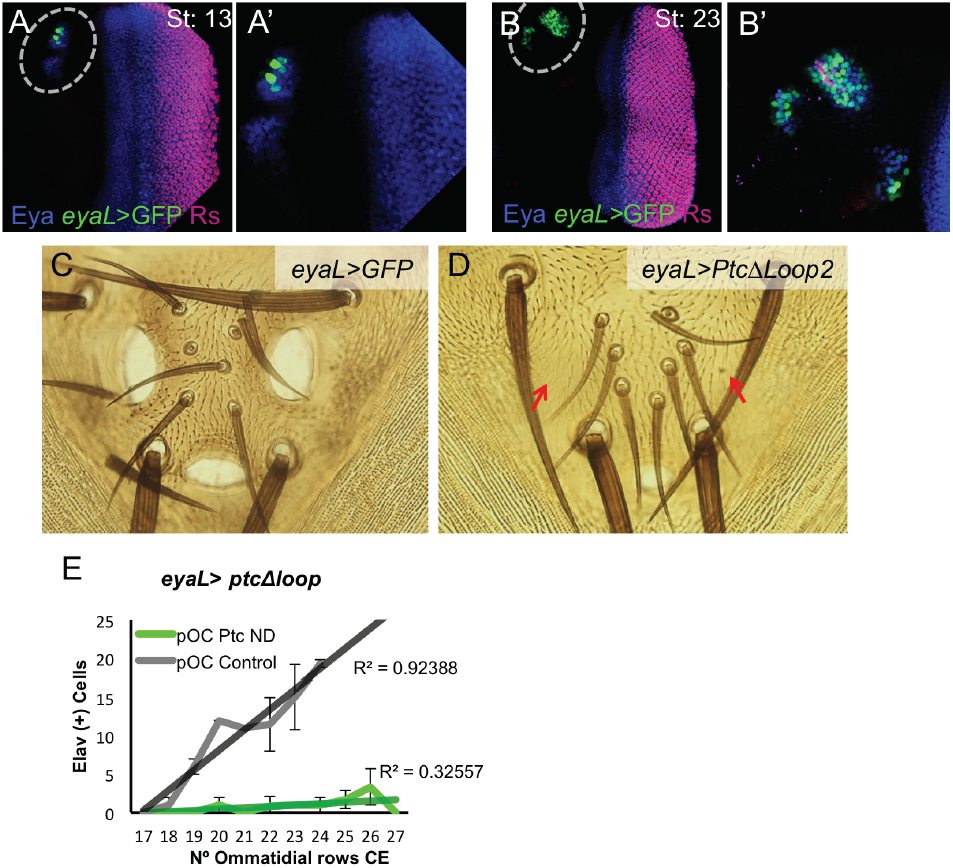
Driving a dominant-negative *Ptc* in the ocelli using the *eyaL-GAL4* blocks ocellar development. (A,B) Enhancer activity of the FlyLight R20D09 GAL4 driver line (“eyaL-GAL4”) before (A: stage 13; A’:close-up) and after (B: stage 23; B’:close-up) R cell differentiation onset. Discs of eyaL-GAL4; UAS-GFP *(eyaL>GFP)* larvae were stained for Eya, GFP and Elav (R cells). Ocellar region is enclosed in the dashed oval. In (A), GFP signal is activated in the Eya-expressing ocellar domains. In later stages (B) GFP signal overlaps Eya. Therefore, EyaL-GAL4 drives expression in the ocellar *eya* domains exclusively. (C,D) Ocellar regions of adult flies. Control (C: *eyaL-GAL4; UAS-GFP*, “eyaL>GFP”) and *eyaL-GAL4; UAS-ptcΔloop2* (D: “eyaL>ptcΔloop2”). PtcΔloop2 acts as a Hh-dominant negative protein (see Material and Methods and references). In *eyaL>ptcΔloop2* flies the ocelli are severely reduced or absent (red arrows). (E) Dynamics of R cells differentiation in the posterior ocellus (pOC) (as Elav-expressing cells) in *eyaL>GFP* (“control”) and *eyaL>ptcΔloop2* (“ptcDN”), with linear fits and R^2^ values. Source data for Figure 2-Figure S1 is available as supplementary material (SD_Figure2_S1E).

**Figure 3-Figure S1.**
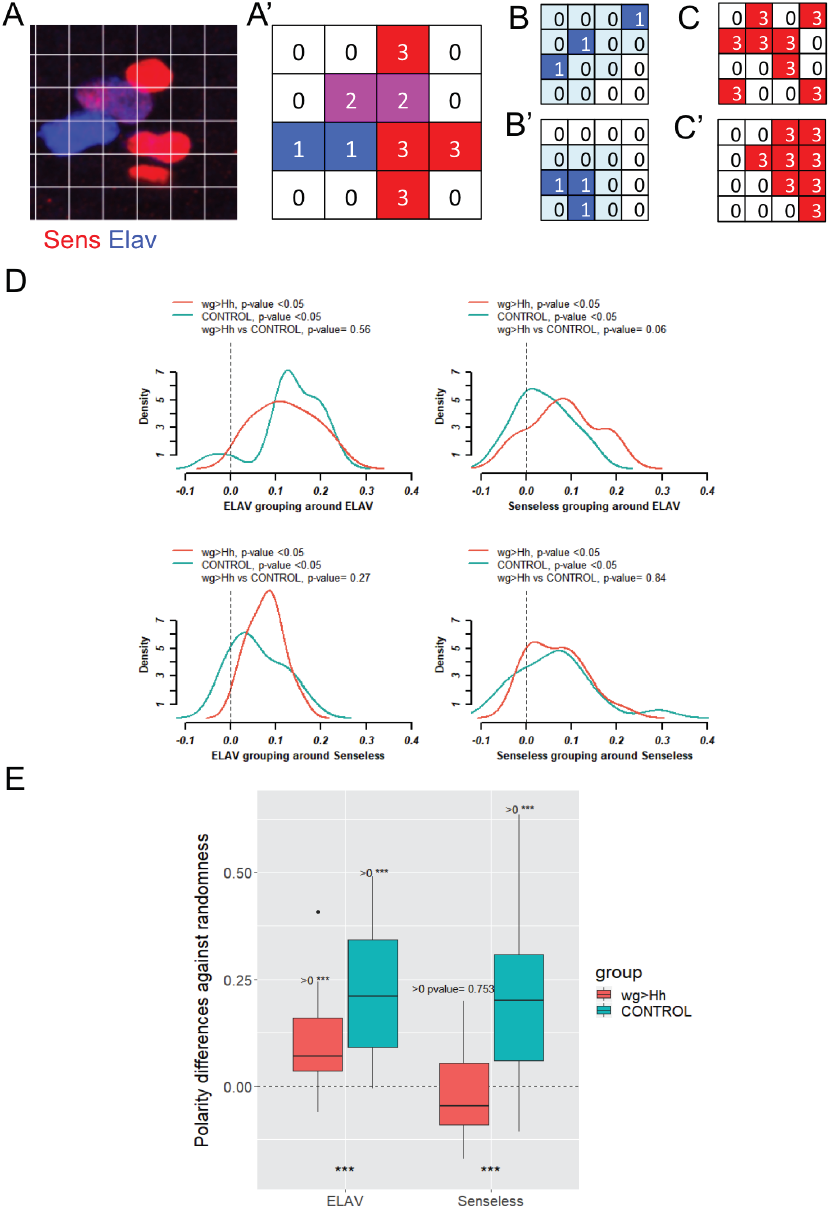
Statistical analysis of Sens/Elav pattern in control and *wg>Hh* ocelli. (A) Image of a Sens and Elav staining with superimposed grid and its translation into a bidimensional matrix (A’), in which the three different states detected, Elav+Sens-, Elav(+)Sens(+) and Elav-Sens+ are coded as 1, 2 and 3, respectively. (B-C’) Examples illustrating the two statistics used to analyze the pattern of Sens and Elav expression. (B,B’) Example of “random” (B) and “grouped” (B’) “1” matrix. Neighbors are marked in light colors. From left to right, the neighbor proportion is 0, 1/8 and 1/5, 0.1 on average for (B), and 2/5, 2/8 and 2/5, 0.4 on average for (B’). (C,C’) Example of “non-polarized” (C) and “polarized” (C’) “3” matrix. Polarity is calculated as the probability of finding a “3” in the last column, estimated using column number as predictor, minus the expected probability of success in the whole matrix (8/16). Polarity will be close to 0 for (C, “non-polarized”) and significantly larger than 0 for (C’, “polarized”). See methods and Results for further details. (D) Represents the departure from random grouping of different cell states (“order”). Both, control and *wg>Hh* patterns show significant ordered grouping for all four comparisons (p<0.05), although they do not differ significantly among them. (E) When the ordered distribution of Elav or Sens along the proximodistal axis (“polarity”) is computed, the pattern in control ocelli is significantly polarized and much more so than in *wg>Hh* samples. Only posterior ocelli were analyzed. Source data for this figure can be found as supplementary material (SD_Figure3_S1).

**Figure 4-Figure S1.**
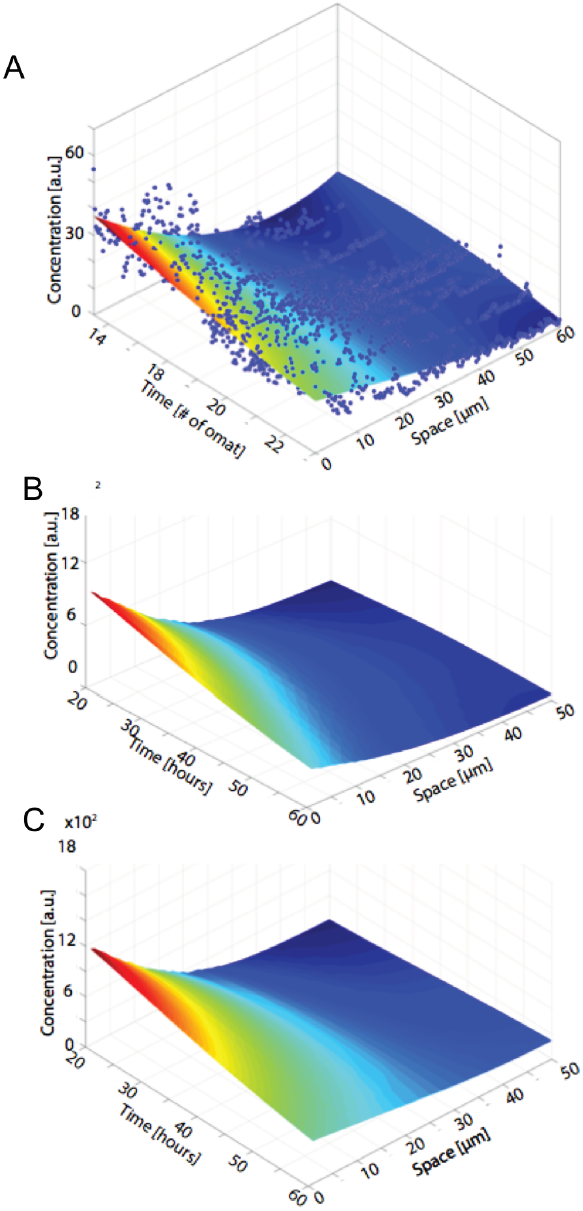
Hh gradient dynamics. (a) Plot of Hh:GFP signal (“concentration” in arbitrary units [a.u.]) as a function of time and space (in μm), obtained from fixed samples at specific developmental time points (as no. of ommatidia in the compound eye)(See supplemental Source Data SD_FigureS3). Model parameters were constrained using this data. (b,c) Plots of Hh dynamics from the simulations, not including (b) or including (c) the attenuation of Ptc expression in differentiating R cells. Note that in (c) (but not in (b)) the Hh gradient spreads farther with time, as observed in the measured profiles (a). Source data for Figure 4-Figure S1A (SD_Figure4_S1) and source code for 3D graph and polynomial adjustment (SD_Figure4_S1Dg_polynadjust) are available as supplementary material.

**Figure 4- Supplementary Table.**
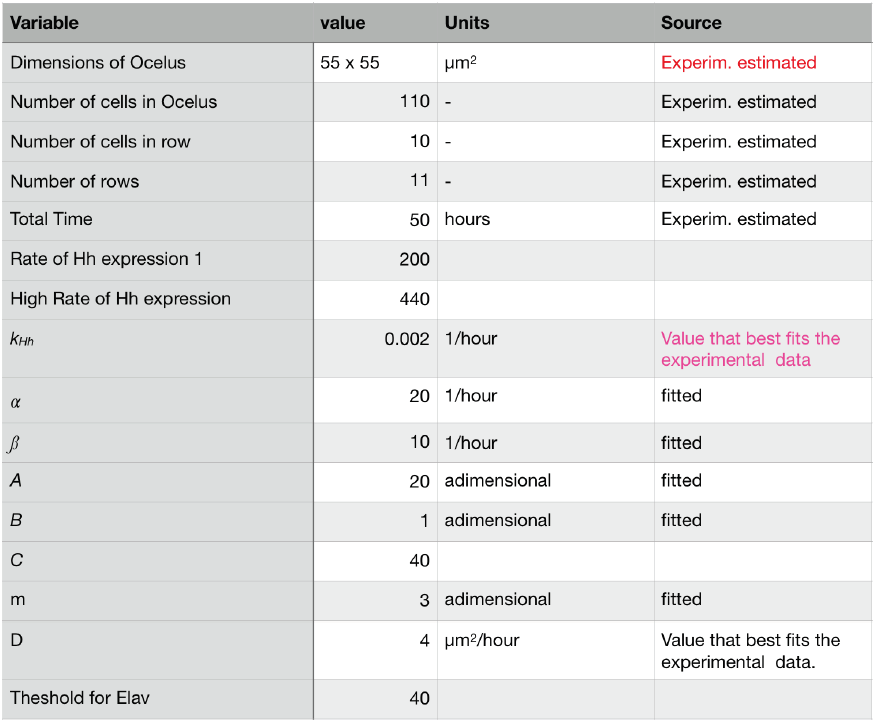
Table of values, units and sources of model parameters.

**Figure 4- Figure S2.**
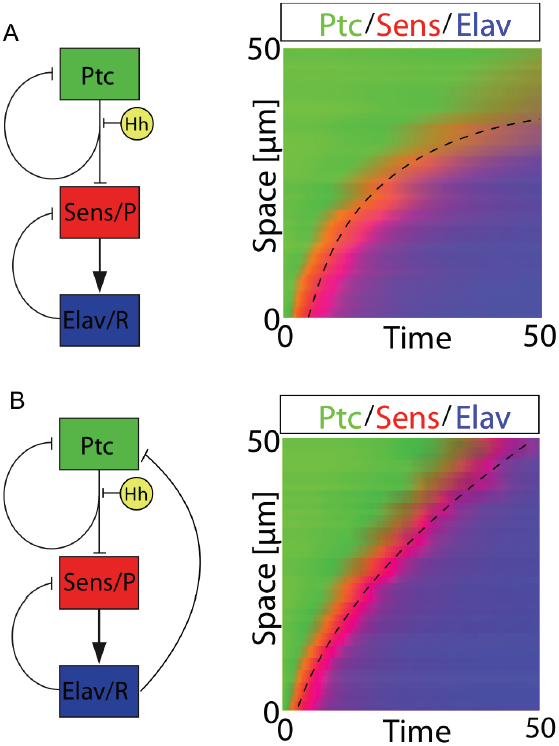
Differentiation dynamics and Ptc attenuation with a constant Hh production rate. Cartoon diagrams of the model for the Hh signaling pathway without considering (A) or considering (B) a negative feedback from Elav-expressing R cells to Ptc and its downstream effects (left), and corresponding spatio-temporal dynamics (right). In (A) R cells (blue) accumulate hyperbolically and do not reach the end of the competent region within the time frame of 50h. In (B) (with negative feedback, all other parameters being the same) R accumulation dynamics is close to linear and R differentiation reaches the end of the competent region. Simulations performed maintaining a constant Hh production rate.

**Figure 4-Figure S3.**
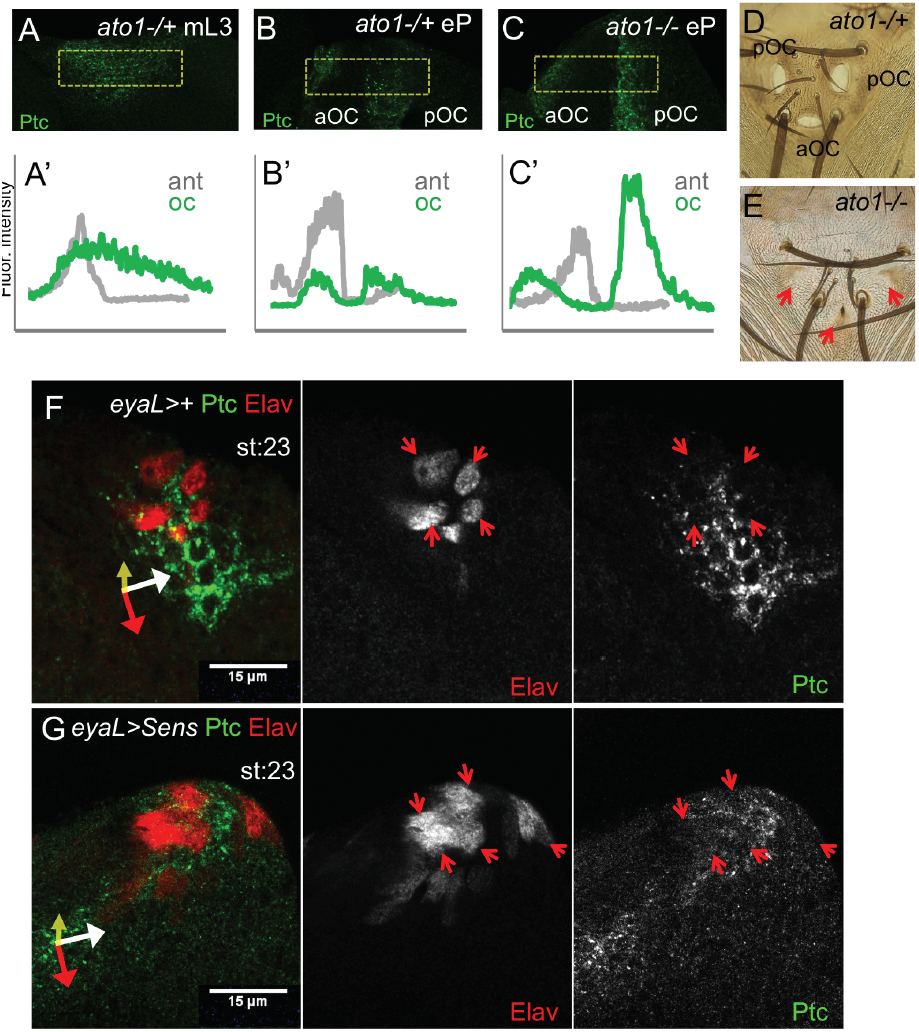
Ptc signal and R cell differentiation. (A-C) Ptc signal in the ocellar regions of mid L3 (mL3; A) and early pupa (eP; B) *ato1+/-* discs, and eP of an *ato1-/-* disc (C). (A’-C’) quantification of Ptc signal in the ocellar regions (“oc”, green) relative to that in the antenna of the same disc (“ant”, grey), this latter used as an internal normalization. In *ato1-/+* controls, the ocellar Ptc signal is similar to that of the antenna (A’; n=6) in early discs but drops in later stages (early pupa: B’, n=5). However, in late stage *ato1-/-* the Ptc signal ratio remains high (C’; n=6). Posterior and anterior ocelli (pOC and aOC) are marked in (B,C). In (A) the split of the Ptc domain in the two ocellar primordia has not yet occurred. (D,E) Adult ocellar complexes of *ato1+/- and ato1-/-* flies. In homozygous *ato1* individuals ocelli fail to develop. (F,G) Control (F: *eyaL>+*) and Sens-expressing (G: *eyaL>Sens*) pOC at st:23 stained for Elav and Ptc. In *eyaL>Sens* there is an increase in the number of Elav cells. Ptc signal is reduced in all Elav cells and, as a consequence, in *eyaL>Sens* Ptc levels are globally reduced also. Red arrows point to Elav cells.

**Figure 4-Supplementary Videos 1 and 2. Time lapse movies of the simulation without (Sup. Video 1) and with (Sup. Video 2) Ptc ngative feedback regulation**.

Upper panel shown the concentration of Hh across the domain. Lower panel shows the cellular concentration of Ptc (green), Sens (red) and Elav (blue). Despite the variable response to Hh due to cell variability, a wave in photoreceptor differentiation (blue cells) can be observed as traveling away from the Hh source. The first movie (no Ptc downregulation) shows that the wave velocity diminishes and stops before reaching the end of the domain. The Hh gradient does not flatten. The second movie includes Ptc negative feedback, and shows how the differentiation wave moves at constant speed and reaches the end of the domain. The Hh gradient flattens.

**Figure 5-Figure S1.**
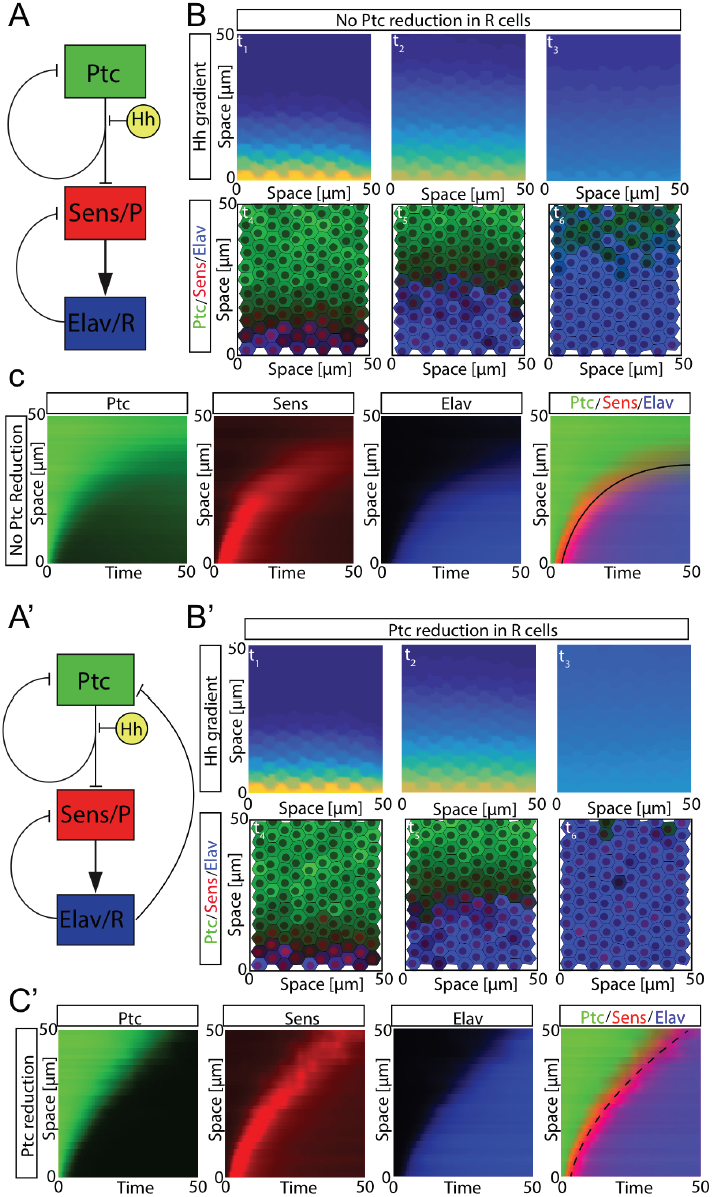
Dynamics of Hh signaling in response to its gradient. (A,A’) Shape of the interactions taken into account in the model. (B,B’) Snapshots of the simulation at different times. Images above depict the Hh profile, while the bottom images represent the cell differentiation state. Red: Sens cells; Blue: Elav R cells; Green: free Ptc (See supp. videos 1 and 2). (C,C’) Space-time plots of free Ptc (green), Sens (red) and Elav (blue) and superposition of the three across the ocellus. The black lines (solid in (C) and dashed in (C’)) are used as a guide to the eye to show the speed of the differentiation wave. Source code for Hh signaling model available as supplementary material (SD_Figure4_script_Hhpathwaymodel).

## METHODS

### Drosophila strains and genetic manipulations

Hh:GFP (BAC) was used to monitor the expression for Hh protein[12]. *ato*^*1*^ is an *atonal* mutant allele (Flybase), and *ato:GFP* is described in [25]. GAL4/UAS crosses were set up at 29°C to maximize GAL4-driven expression, except when indicated. The *hh-GAL4, UAS-GFP:Hh* strain was used as reporter for Hh expression[26]. Elav-Gal4 (Flybase) was used to drive UAS-*H2B-mCherry-P2A-eGFP-PH* line[27] in differentiated R cells, allowing the distinction between nuclei (mCherry) and cell membranes (eGFP) (experiment at 25°C). The FlyLight [28] GAL4 line R20D09 from *eya* (herein referred as EyaL-GAL4) was used to drive UAS transgenes specifically in the anterior and posterior ocellar competence domains (ED1). GMRtdTom was used as a reporter of Glass to monitor the PRs cells and membranes [29]. UAS lines used were: UAS-nlsGFP (Flybase), UAS-Ci^FL^ [14] and UAS-GFP-*ptc*Δloop2 (UAS-*ptc*DN)[2]. Quantification of number of R cells over-time was performed in the wild-type strain Oregon-R at 25°C. To perturb the normal distribution of Hh, a GFP-tagged Hh (UAS-GFP:Hh [20]) was driven with the *wg2.11-GAL4* strain *(wg2.11-GAL4; UAS-GFP:Hh*, or “wg>Hh”). wg2.11 is an enhancer of the *wg* gene that is expressed surrounding the ocellar region and overlapping the prospective interocellar region in the eye imaginal disc (described in [15] and see results).

### Immunofluorescence

Medium-late third instar larvae and pupae were dissected and fixed according to standard protocols. Immunostainings were performed as previously described [30]. We used the following primary antibodies: rabbit anti-GFP at 1:1000 (Molecular Probes), rat anti-RFP at 1/500 (Chromotek), rabbit anti-βGal at 1:1000 (Cappel). Mouse anti-Eya 10H6 at 1:400, rat anti-Elav 7EBA10 at 1:1000 and mouse anti-Ptc at 1:100 were from the Developmental Studies Hybridoma Bank, University of Iowa (DSHB, http://dshb.biology.uiowa.edu). Aliquots of mouse anti-Sens at 1:250 were gifts from Andrew Jarman (the University of Edinburgh), Bassem Hassan (ICM, Paris), Rosa Barrio (Biogune, Leioa) and Xavier Franch (IBE-UPF, Barcelona), and rat anti-Ci 2A1 at 1:5 was a gift from Bob Holmgren (Northwestern University). Imaging was carried out on Leica SP2, SPE or SP5 confocal set-ups.

### Measurement of the Hh:GFP signaling gradient dynamics

Eye discs from the BAC Hh:GFP strain were dissected from 96-130 hours after egg laying (grown at 25°C) and stained simultaneously. Number of discs per experiment was >10 and one representative example was shown. Developmental stage was determined as number of ommatidial rows in the region of the compound eye. Imaging was carried out in a Leica SP5 confocal setup with the same settings. Lasers were previously warmed up during 1h. Fluorescence Intensity measurements were obtained with Fiji [31] by selecting a ROI across the ocellar complex. Then a Plot Profile was generated for the ROI and the quantitative data obtained were processed in Excel.

#### R cell recruitment over-time

Medium-late third instar OR-R larvae and pupae were dissected and stained with anti-Elav to monitor the degree of differentiation from the stage 17 ommatidia to stage 27 ommatidia. The total number of samples quantified was 83 for both ocelli. Samples per time point ranged from five to 12. To analyze the correlation of the number of ocellar photoreceptors cells (Rs) and developmental time, as measured by the number of ommatidia rows in the compound eye, we performed an univariate linear regression, using the formula:

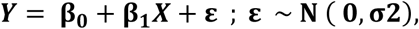

where Y is the number of R cells; X is the number of ommatidial rows in the compound eye; β_0_ is the intercept coefficient; β_1_ is the number of ommatidial row coefficient and ε is the regression error. The model was estimated by the least squares method using *lm()* function in *Rsoftware* and validated checking for normality, independence and homoscedasticity of residuals. The analysis shows a statistically significant linear dependence between PR cell number and developmental time, either when considering PR cell number of the anterior or the posterior ocelli individually or aggregating the data from both ocelli. The table below summarizes the statistics results of the linear regression. Source data for Figure 2D is available as supplementary material (SD_Figure2_A’_D).

|  | pOC | aOC | pOC+aOC |
| --- | --- | --- | --- |
| *Coef. Estimated* | 2.8067 | 1.039 | 3.8386 |
| *Coef. p-value* | <2e-16 | 2.39E-15 | <2e-16 |
| *R-squared* | 0.7722 | 0.5995 | 0.7794 |
| *Adjusted R-squared* | 0.7689 | 0.5937 | 0.7762 |
| *Model p-value* | 2.2E-16 | 2.391E-15 | < 2.2e-16 |

#### Quantification of adult ocelli R cell number

Brain preparations, with the ocelli attached, were dissected from newly hatched (0-1 days) adults and stained with anti-Elav and counterstained with Rhodamine-phalloidin (cell membranes) and DAPI (nuclei). Ocelli were imaged as z-stacks on an SPE Leica confocal setup and reconstructed using Imaris (Bitplane) for quantification.

#### Spatial statistics of Elav and Sens pattern under normal and perturbed Hh distribution

We imaged as confocal z-stacks ocellar regions stained for Sens and Elav from control (Oregon-R strain; N=19) or *wg2.11>GFP:Hh* (N=18), in the range of 18-23 ommatidia stage. Three cell states can be observed: 1: [Sens-, Elav+], 2: [Sens(weak), Elav(weak)] and 3: [Sens+, Elav-] that correspond to differentiating photoreceptors, the transition between precursors and photoreceptors, and precursors, respectively. To obtain a bidimensional description of the distribution of these cells types in the tissue, we superimposed an orthogonal grid (ImageJ: Analyze>Tools>Grid) on a maximal projection of the z-stack sections comprising all Sens and Elav signals. The grid’s cell size is set to correspond approximately to the size of a cell’s nucleus, so that, in general there is only one nucleus per grid’s cell. When a nucleus spans two or more cells in the grid, its position is allocated to the grid’s cell where most of the signal is. Then, a 1, 2 or 3 is assigned to each grid cell according to its Sens and Elav signal. A grid cell with no signal is assigned a “0”. The result is a two-dimension matrix of positions of the three states per sample (Figure 3_Figure S1).

### Statistical analysis of Elav and Sens expression patterns

In order to test the departure from a random pattern of Sens and Elav expression we defined two statistics: “grouping” and “polarity”. Importantly, the degree of polarization will tell whether the pattern is compatible with a wave-like organization. For the analysis, each matrix comprising 1, 2 and 3 cell types (Elav+Sens-, Elav+Sens+ and Elav-Sens+, respectively) is split into two matrices, one in which 2 is identified as 1 and another in which 2 is identified as 3, since “2” is expression of 1 and 3 in the same cell. This allows a straightforward statistical analysis.

“Grouping” is defined as the departure from a random proportion of neighbors of a given type for each cell expressing Elav or Sens. For each cell *i*, the proportion of Elav or Sens-expressing neighbors, *p*_*i*_ is calculated as

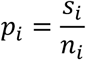

where *s*_*i*_ is the number of neighbors of a given type and *n*_*i*_ is the total number of neighbors (note that this number will depend on the position of the cell within the matrix, with cells in the center with more neighbors (8) than if in the periphery). Grouping is a global property of the ocellus so the estimation of grouping for the whole ocellus could be reduce to count the number of Elav or Sens expressing cells relative to total cells in the neighborhood:

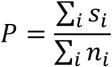

However, this grouping is strictly dependent on the proportion of Elav or Sens in the ocellus, so in order to obtain an unbiased measure of grouping the total proportion of cells expressing a factor needs to be substracted from the proportion of this factor in the neighborhood. As a correction of the statistic thus defined we actually consider the total proportion as:

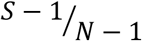

Where S is the total number of cells expressing the factor in the ocellus and N the total number of cells in it. We have to subtract 1 from the numerator and denominator because each time we calculate the proportion of neighbors, we focus non-randomly on a cell expressing the factor (effectively we are “removing” one case from the sample) so the proportion of success in the neighborhood that can be expected in a random matrix would be lower than the actual proportion. Then, grouping of cells of the same type is expressed as follows:

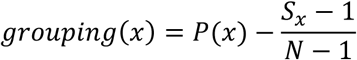

Where *x* is the expressed factor, Elav or Sens.

However, if we consider grouping of Elav around Sens or Sens around Elav making the previous correction is not needed, because this time the expected proportion of success in the neighborhood in a random matrix coincides with the total proportion in the ocellus, so for the case of grouping of cells of one type *(y)* around a cell of the other type *(x)*, the grouping would be defined as:

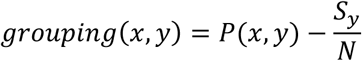

Where *x* is the expressed factor in a cell, and *y* is the other factor, expressed in the cell’s neighborhood.

“Polarity” measures the ordered succession of cells states along a spatial axis. In our case, it is the “proximodistal” axis with “proximal” defined as the position closest to the endogenous Hh source. For each matrix and each factor it is possible to define a dichotomous response variable *Y* which classifies a cell as expressing a factor, 1, or not, 0. So we can define a logistic regression model to predict the expression of this factor in a cell using column position, *X*, as predictor:

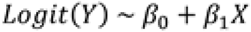

The hypothesis for Elav is that its expression will be “proximal”, that is the left or first column, whereas Sens will be “distal”, that is the right or last column, so after the estimation of the model for each factor we will use these models to predict the probability of finding ELAV at the first column and Senseless at the last one. The following expression defines them:

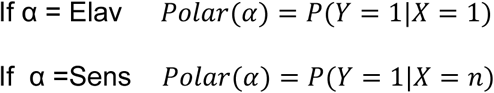

where *n* is number of columns in the matrix and α the factor used.

This probability has to be compared with the probability of finding expression of α at that column randomly or what is the same with no predictor used, which coincides again with the total proportion of the factor in the matrix. Polarity is then defined as follows:

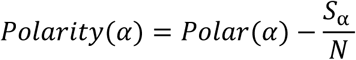

Where *S*_α_ is the number of cells expressing the factor and *N* the number of cells in the matrix.

Groups comparison: For each matrix 4 measures of “grouping” and 2 of “polarity” were estimated:

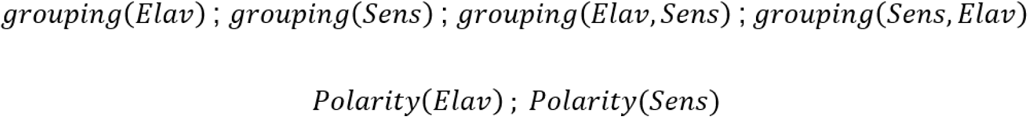

Then they were calculated for every matrix and plotted (Figure 3-Figure S1). In order to test for significant grouping differences between control and *wg>Hh*, a Welch’s test for unequal variances was performed for each grouping variable:

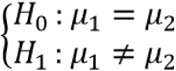

Since we aimed at testing if there was a pattern of Elav and Sens expression, we had to check that they were not distributed randomly, so grouping should be larger than 0. A Student‘s t-test for each grouping distribution and experimental group, control or *wg>Hh*, was performed:

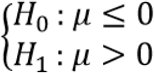

The same hypothesis was posed and the same test performed for polarity, first to check if these groups were significantly different from one another and then to check if the polarity was larger than 0.

Statistics and data treatment were performed in R software. Data matrices were imported to R from.csv.

### Adult cuticles dissections

The dorsal head capsules were dissected in PBS1X. Brain tissues and proboscis were removed from the samples. All the structures were incubated overnight in Hoyer’s:Lactic Acid (1:1) solution at 80°C [32]. Imaging was carried out on a Leica DM500B microscope with a Leica DFC490 digital camera. All images were processed with Fiji [31].

### Modeling the Hh pathway in the *Drosophila* ocelli

Simulations were performed using an in-house computational script developed in Matlab^®^ (The Mathworks). This script is available as source code (SD_Figure4_script_Hhpathwaymodel). Equations are discretized in space and time using an Euler approach, with adimensional concentrations but dimensional variables in space and time. The model is based on a hybrid approach that combines partial differential equations (PDEs) solved in a continuous space and ordinary differential equations (ODEs) that are solved in a discrete space. The PDEs account for the diffusive extracellular signals, while the ODEs account of the intracellular reactions. Cells are simulated as two-dimensional regions in a hexagonal Voronoi diagram, with cell-to-cell variability introduced as gamma-distributed values for each of the kinetic constants of the reactions involved, with a standard deviation of 20% of the mean value.

The equations define a simplified representation of the Hh signaling pathway, illustrated in Figure 3. The set of interactions that the model takes into account are the following:

#### Diffusion of the Hedgehog (Hh) morphogen

Hh is secreted by producing cells in the intervening region between the anterior and posterior ocellar competent regions (“Hh source”) and then disperses generating a concentration gradient. The mechanism by which Hh disperses is not totally understood, and several studies propose that Hh travels through cytonemes[33] as an alternative to diffusion. Overall, the highly noisy spatiotemporal profile of Hh distribution in the ocellus (Figure 4-Figure S1A) can be fitted with a polynomial that decreases non-linearly when moving away from the Hh source. Our model simplifies the details of Hh transport as two-dimensional diffusion. This approach successfully reproduces the experimental data of shape and dynamics of the Hh profile (see Figure S3). The equation that governs Hh dynamics is:

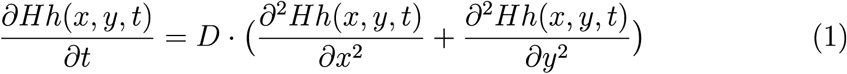

Our model approximates the Hh source as a continuous supply of Hh at one of the boundaries of the ocellus. The experimental data shows that Hh expression by the Hh-producing cells cells, monitored by a Hh:GFP BAC, gradually decreases to 50% of its initial values during the period through which cell differentiation is taking place. This is introduced in our model as a continuous reduction in the Hh production rate at the production boundary to around 50% of its initial value. However, similar computational results are obtained if the Hh production rate is maintained constant (see Figure 4-Figure S2). Source data for Hh:GFP profile quantification is available as supplementary information (SD_Figure4_S3A‘_B‘).

#### Binding of Hh to its receptor Ptc

Hh binds to its receptor Ptc irreversibly to form a complex (*Ptc-Hh*)[19–21], following the scheme:

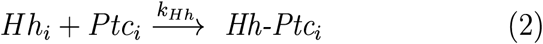

Where *k*_*Hh*_ corresponds to the affinity rate constant of the interaction. *Hh_i_* corresponds to the amount of Hh that a given cell *i* is receiving, computed at each time step as the average value of Hh over the whole cell area of cell *i*. In this way, the continuous value of Hh computed in Eq. 1 is converted to a discrete value for each cell in the population *Hh*_*i*_. This value is then used to compute the amount of Hh that binds to Ptc via Eq. 2 as an ODE that is solved for each cell in the hexagonal lattice:

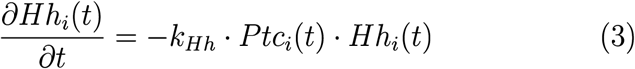

This ODE equation is then solved continuously in time but discretely in space, i.e., for each cell *i* in the population. Then, the amount of Hh molecules consumed by each cell *i* in each particular position is subtracted from the continuous spatial variable Hh in the corresponding position. The resulting Hh profile is then computed at the next time step via Eq. 1.

#### Expression of Ptc and binding to Hh

The amount of Hh that reaches a given cell in the population interacts with the free form of its receptor, Ptc. In the absence of Hh, free Ptc acts, indirectly through inhibition of the signal transducer Smo, as a repressor of Hh signaling target genes. This repression is set in the model as sigmoidal function of Ptc, with cooperativity *m*=3 (slightly higher or lower values of *m* also reproduce the experimental results). Since one of Hh targets is Ptc itself, the sigmoidal repression is introduced in the equation corresponding to Ptc, forming a direct negative feedback loop. In addition, a constant degradation of Ptc is introduced to ensure a dynamic equilibrium in its concentration. Taking this into account, the dynamics of Ptc is described by the following ODE:

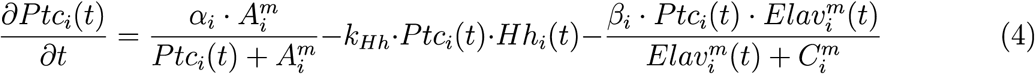

where ☐ and *ß* corresponds to the rate constant for production and degradation. *A* corresponds to the half maximal concentration of the sigmoidal curve, and *m* sets the slope of the sigmoidal. The next term accounts for the binding of Ptc and Hh, following Eq 2.

The second version of the model includes a reduction of available (“free”) Ptc in terminally differentiated photoreceptors. This is simplified in the model by adding the last term in Eq. 4 in the form of a Hill function dependent on *Elav*, a marker of photoreceptor (“R”) fate.

#### Expression of Senseless (Sens)

One of the relevant Hh signaling pathway targets (albeit likely indirect) is *senseless* (Sens), a Zn-finger transcription factor required for ocellar photoreceptor differentiation downstream of the proneural gene *atonal* [10, 34]. Our model described the dynamics of expression of Sens by the following ODE:

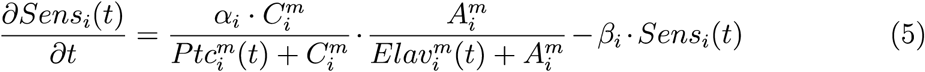

where the expression of *Sens* is mediated by simple direct repression by *Ptc*, where the half maximal concentration of the sigmoidal correspond to *B*. In addition, we have observed that during ocellar differentiation *Sens* expression is also lost in terminally differentiated photoreceptors. We represent this loss of *Sens* expression in the models as a direct repression by *Elav* in each cell *i*. To make this repression stronger than the repression by Ptc, the second term is elevated again to *m*.

#### Differentiation into a terminal photoreceptor cell

The events downstream of Sens that result in a terminally differentiated photoreceptor cell are also simplified in a single activation of the Elav gene. Its expression is assumed as directly proportional to the amount of Sens, with a sigmoidal degradation of Elav. Therefore, the equation for the dynamics of Elav takes the form:

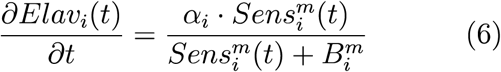

Once the concentration of Elav reaches a given threshold value in a cell *i*, the model assumes an irreversible transition to a differentiated photoreceptor.

